# Integrative phylogenetic analysis of the genus *Episoriculus* (Mammalia: Eulipotyphla: Soricidae)

**DOI:** 10.1101/2024.02.15.580476

**Authors:** Yingxun Liu, Xuming Wang, Tao Wan, Rui Liao, Chen Shunde, Shaoying Liu, Bisong Yue

## Abstract

Shrews in the genus *Episoriculus* are among the least-known mammals in China, where representatives occur mainly in the Himalayan and Hengduan mountains. We sequence one mitochondrial and three nuclear genes from 77 individuals referable to this genus, collect morphometric data for five shape and 11 skull measurements from 56 specimens, and use museum collections and GenBank sequences to analyze phylogenetic relationships between this and related genera in an integrated molecular and morphometric approach. Whereas historically anywhere from two to eight species have been recognized in this genus, we conclude that six (*E. baileyi*, *E. caudatus*, *E. leucops*, *E. macrurus*, *E. sacratus*, *E. soluensis*) are valid. We dissent from recent systematic reviews of this genus and regard *E. sacratus* to be a valid taxon, *E. umbrinus* to be a subspecies of *E. caudatus*, and transfer *E. fumidus* to *Pseudosoriculus*. Our record of *E. soluensis* is the first for China, and expands the previously recognized distribution of this taxon from Nepal and NE India into the adjacent Yadong and Nyalam counties. One further undescribed *Episoriculus* taxon may exist in Tibet.

## Introduction

The genus *Episoriculus*, originally established as a subgenus of *Soriculus*, occurs throughout southwest China, India, Nepal, and Vietnam [1–9]. This subgenus was relegated to full generic status by Repenning [10] on grounds of significant differences in tooth morphology from other species of *Chodsigoa* and *Soriculus*—a taxonomy followed by Jameson & Jones [11], Hutterer [12], Wilson & Reeder [13], and Wilson & Mittermeier [3].

The number of valid species of *Episoriculus* has been the subject of debate, with 2–8 species recognized (Table 1). Allen [14] described *S. macrurus*, *S. caudatus sacratus*, and *S. caudatus umbrinus*. Ellerman & Morrison-Scott [1] proposed *Episoriculu*s as a subgenus of *Soriculus*, and included *S. leucops* and *S. caudatus* (with subspecies *S. c. caudatus*, *S. c. baileyi*, *S. c. fumidus*, *S. c. sacratus*, and *S. c. umbrinus*). Honacki *et al.* [6] proposed that *Episoriculus* included four species, and considered *S. baileyi* and *S. fumidus* to be valid taxa. Hoffmann [7] similarly recognized four species, although these were not entirely consistent with those of Honacki *et al.* [6], for *S. baileyi* was relegated to a subspecies of *S*. (*E*.) *leucops*, and *S. (E.) macrurus* was placed in this genus. Corbet & Hill [8], and Wilson & Reeder [9, 13] followed this arrangement. Motokawa and Lin [15] elevated *S. baileyi* to full species based on morphology. Based on the karyotypes and differences in skull morphology, Motokawa *et al*. [16] considered that *E. caudatus* should be divided into the larger *E. caudatus* and smaller *E. sacratus* (with subspecies *E. s. soluensis* from Nepal and Sikkim, *E. s. umbrinus* from Assam, Myanmar, and Yunnan, China, and *E. s. sacratus* from Sichuan, China). He *et al.* [15] noted that *P. fumidus* did not belong to *Episoriculus*. Based on *CYTB* gene, Abramov *et al.* [2] promoted *E. soluensis* to full species, assigned *E. fumidus* to a new genus *Pseudosoriculus*, and arranged seven species (*E. baileyi*, *E. caudatus*, *E. leucops*, *E. macrurus*, *E. sacratus*, *E. soluensis*, and *E. umbrinus*) in *Episoriculus*. Wilson & Mittermeier [3] recognized eight species, including *P. fumidus*.

**Table 1.**
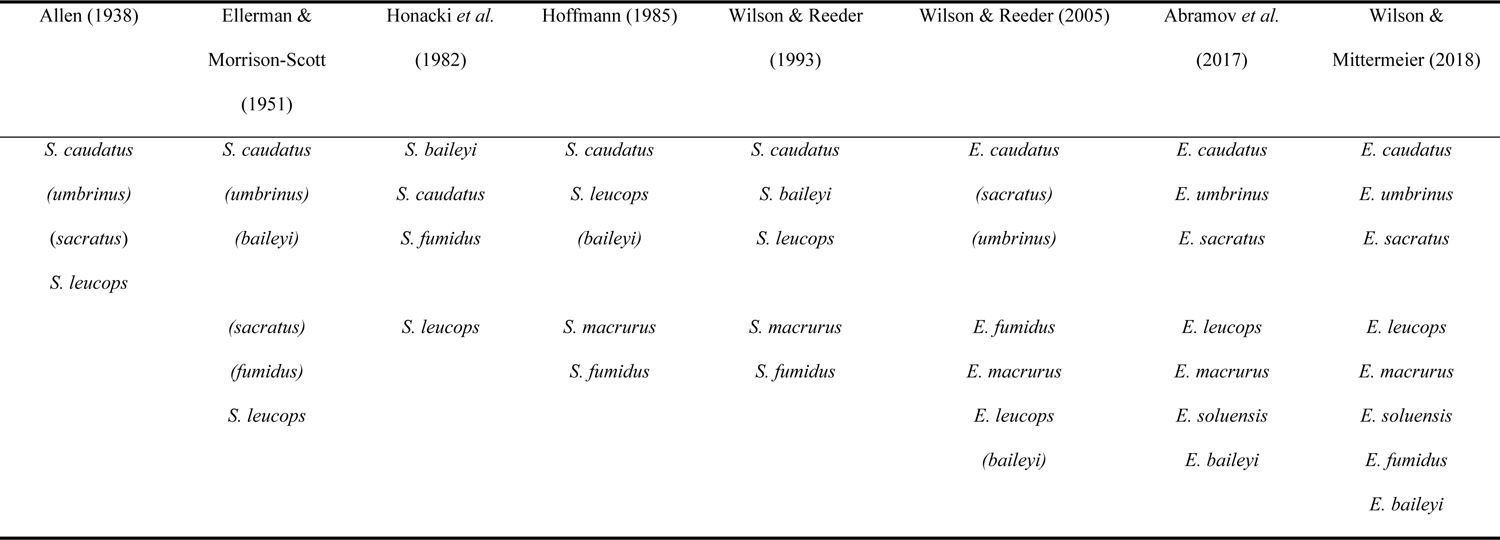
Major classification systems of the genus *Episoriculus*. The name of the species in parentheses indicates that the species is a subspecies of the previous species.

Throughout these various classifications the taxonomic status of *E. caudatus*, *E. leucops*, and *E. macrurus* has been relatively stable, but the taxonomy of *P. fumidus, E. sacratus*, *E. umbrinus*, *E. baileyi*, and *E. soluensis* has not. We report new molecular data and morphological comparisons in an integrated phylogenetic and morphological analysis to clarify the taxonomic status of species in the genus *Episoriculus*.

## Materials and methods

### Ethics statement

All specimens were collected in accordance with regulations in China for implementation of the protection of terrestrial wild animals (State Council Decree [1992] No. 13). Collecting protocols and the research project were approved by the Ethics Committee of Sichuan Academy of Forestry (no specific permit number).

### Sampling and sequencing

Of 77 specimens collected from within China, 31 were attributed to *E. macrurus*, 18 to *E. caudatus*, 10 to *E. leucops*, 5 to *E. umbrinus*, 4 of each to *E. sacratus* and *E. soluensis*, and 2 to *E.* sp. (Table 2, Fig. 1) following Wilson & Mittermeier [3], Hoffmann [7], and Smith & Xie [14]. Of recognized species, we had no specimens of *E. baileyi*. Voucher specimens of animals reported on in this study have been deposited in collections of the Sichuan Academy of Forestry. Muscle and liver tissue of all specimens were preserved in 95% ethanol, then stored at −75°C until DNA extraction.

**Fig. 1.**
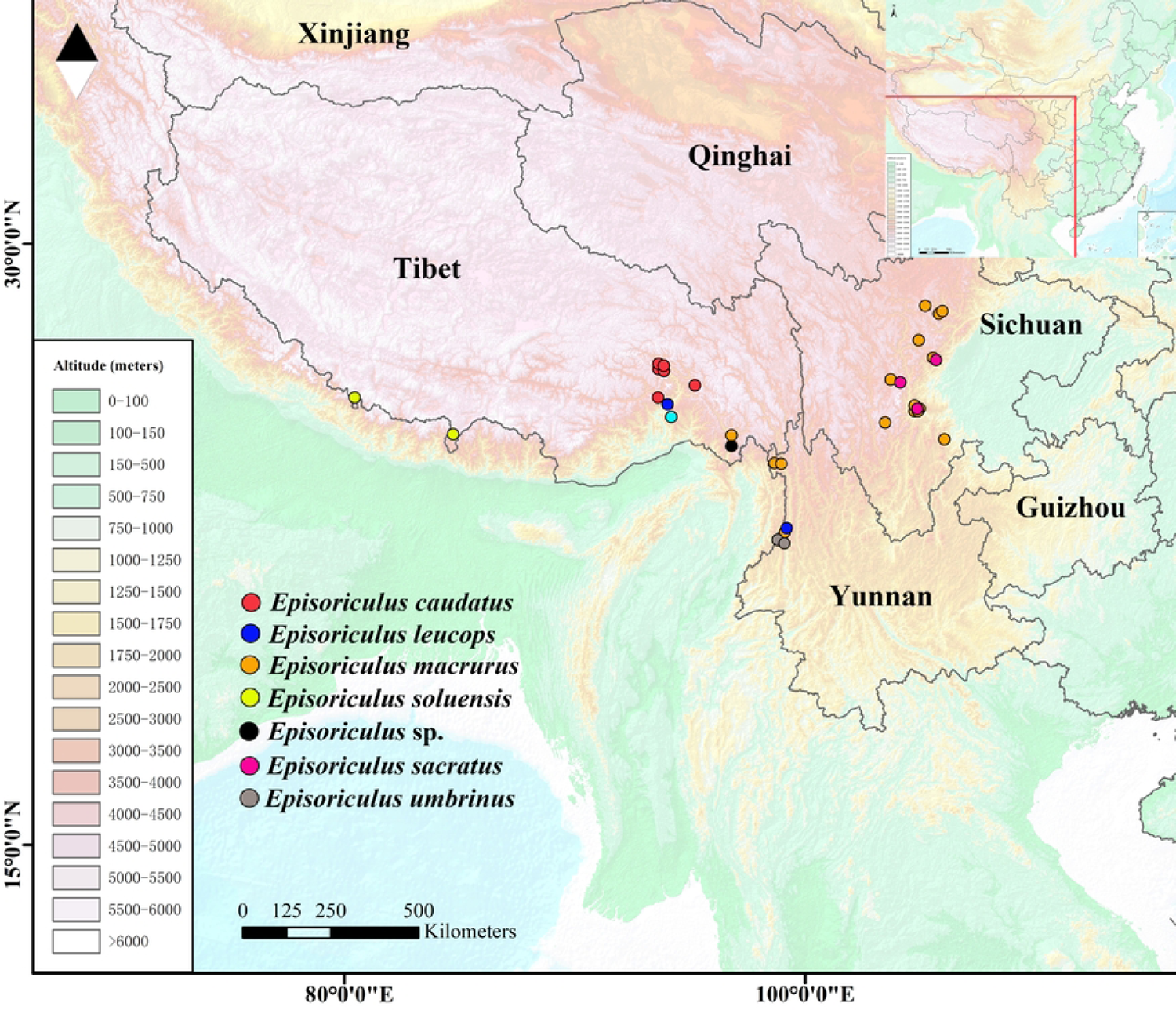
Map of genus *Episoriculus*, showing localities sampled for this study. This is the Fig. 1 legend.

**Table 2.** Samples and sequences of *Episoriculus* used for molecular analyses.

| Species | Species ID | Sequence No. | Locality | longitude | Latitude | Altitude(m) | *CYTB* | Genbank accession No. |  |  |
| --- | --- | --- | --- | --- | --- | --- | --- | --- | --- | --- |
|  |  |  |  |  |  |  |  | *APOB* | *BRCA1* | *RAG2* |
| *Episoriculus sacratus* | SAF15216 | LS15073 | Lushan, Sichuan | 103.04592 | 30.66332 | 2018 | MK962202 | / | MN032211 | MN032279 |
| *Episoriculus sacratus* | SAF06888 | ELSTNPB01007 | Erlangshan, Sichuan | 102.29916 | 29.88547 | 2450 | MK962204 | MN032147 | MN032213 | MN032281 |
| *Episoriculus sacratus* | SAF06902 | ELSTNPB02004 | Erlangshan, Sichuan | 102.29916 | 29.88547 | 2450 | MK962207 | MN032149 | MN032214 | MN032283 |
| *Episoriculus sacratus* | SAF11594 | BLG012 | Balanggou, Sichuan | 101.98069 | 30.41007 | 2460 | MK962218 | / | MN032223 | / |
| *Episoriculus sacratus* | SAF06934 | ELS260B02003 | Erlangshan, Sichuan | 102.31112 | 29.87314 | 2230 | MK962203 | / | MN032212 | MN032280 |
| *Episoriculus sacratus* | SAF06920 | ELS260A02003 | Erlangshan, Sichuan | 102.30982 | 29.87294 | 2220 | MK962217 | / | MN032222 | MN032292 |
| *Episoriculus caudatus* | SAF13199 | NJ13033 | Lushui, Yunnan | 98.70969 | 25.98614 | 2719 | MK962237 | MN032174 | MN032241 | MN032310 |
| *Episoriculus umbrinus* | SAF13257 | NJ13091 | Gongshan, Yunnan | 98.44691 | 27.77396 | 3320 | MK962198 | / | MN032207 | / |
| *Episoriculus umbrinus* | SAF180044 | YN18021 | Mountain Gaoligong, Yunnan | 98.7111 | 25.97795 | 2570 | MK962223 | MN032163 | MN032228 | MN032297 |
| *Episoriculus caudatus* | SAF07474 | XZRAP05005 | Nyingchi, Tibet | 94.962862 | 30.00965 | 2040 | MK962213 | MN032155 | MN032218 | MN032289 |
| *Episoriculus caudatus* | SAF07492 | XZRAP07004 | Nyingchi, Tibet | 94.9631 | 30.00942 | 2020 | MK962212 | MN032154 | MN032217 | MN032288 |
| *Episoriculus caudatus* | SAF07568 | XZXCXY04003 | Xiachayu, Tibet | 97.01193 | 28.51268 | 1640 | MK962224 | / | MN032229 | MN032298 |
| *Episoriculus caudatus* | SAF07483 | XZRAP06001 | Nyingchi, Tibet | 94.95855 | 30.00766 | 2050 | MK962214 | MN032156 | MN032219 | MN032290 |
| *Episoriculus caudatus* | SAF07566 | XZXCXY04001 | Xiachayu, Tibet | 97.01193 | 28.51268 | 1640 | MK962206 | / | / | / |
| *Episoriculus caudatus* | SAF07502 | XZRAP08004 | Nyingchi, Tibet | 94.95878 | 30.01025 | 2040 | MK962219 | MN032159 | MN032224 | MN032293 |
| *Episoriculus caudatus* | SAF07501 | XZRAP08003 | Nyingchi, Tibet | 94.95878 | 30.01025 | 2040 | MK962222 | MN032162 | MN032227 | MN032296 |
| *Episoriculus caudatus* | SAF07472 | XZRAP05003 | Nyingchi, Tibet | 94.962862 | 30.00965 | 2040 | MK962221 | MN032161 | MN032226 | MN032295 |
| *Episoriculus caudatus* | SAF07490 | XZRAP07002 | Nyingchi, Tibet | 94.9631 | 30.00942 | 2020 | MK962215 | MN032157 | MN032220 | / |
| *Episoriculus caudatus* | SAF07535 | XZLZT02001 | Nyingchi, Tibet | 95.00633 | 30.02289 | 2020 | MK962211 | MN032153 | / | MN032287 |
| *Episoriculus caudatus* | SAF07470 | XZRAP05001 | Nyingchi, Tibet | 94.962862 | 30.00965 | 2040 | MK962216 | MN032158 | MN032221 | MN032291 |
| *Episoriculus* sp. | SAF11247 | MT11162 | Motuo, Tibet | 95.4814 | 29.43014 | 2832 | MK962200 | MN032145 | MN032209 | MN032277 |
| *Episoriculus* sp. | SAF11283 | MT11198 | Motuo, Tibet | 95.0095 | 29.40422 | 3118 | MK962199 | MN032144 | MN032208 | MN032276 |
| *Episoriculus leucops* | SAF180058 | YN18035 | Mountain Gaoligong, Yunnan | 98.7145 | 25.95919 | 2250 | MK962232 | MN032169 | MN032237 | MN032306 |
| *Episoriculus leucops* | SAF180035 | YN18012 | Mountain Gaoligong, Yunnan | 98.7111 | 25.97795 | 2570 | MK962231 | MN032168 |  | MN032305 |
| *Episoriculus leucops* | SAF13190 | NJ13024 | Lushui, Yunnan | 98.68388 | 25.97259 | 3150 | MK962234 | MN032171 | MN032239 | MN032308 |
| *Episoriculus leucops* | SAF180036 | YN18013 | Mountain Gaoligong, Yunnan | 98.7111 | 25.97795 | 2570 | MK962227 | MN032165 | MN032232 | MN032301 |
| *Episoriculus leucops* | SAF180033 | YN18010 | Mountain Gaoligong, Yunnan | 98.7111 | 25.97795 | 2570 | MK962229 | MN032167 | MN032234 | MN032303 |
| *Episoriculus leucops* | SAF180034 | YN18011 | Mountain Gaoligong, Yunnan | 98.7111 | 25.97795 | 2570 | MK962233 | MN032170 | MN032238 | MN032307 |
| *Episoriculus leucops* | SAF180045 | YN18022 | Mountain Gaoligong, Yunnan | 98.7111 | 25.97795 | 2570 | MK962226 | / | MN032231 | MN032300 |
| *Episoriculus leucops* | SAF180040 | YN18017 | Mountain Gaoligong, Yunnan | 98.7111 | 25.97795 | 2570 | MK962228 | MN032166 | MN032233 | MN032302 |
| *Episoriculus leucops* | SAF180037 | YN18014 | Mountain Gaoligong, Yunnan | 98.7111 | 25.97795 | 2570 | MK962230 | / | MN032235 | MN032304 |
| *Episoriculus leucops* | SAF11202 | MT11017 | Motuo, Tibet | 95.126722 | 29.3665 | 2100 | MK962235 | MN032172 | MN032240 | MN032309 |
| *Episoriculus macrurus* | SAF180039 | YN18016 | Mountain Gaoligong, Yunnan | 98.7111 | 25.97795 | 2570 | MK962166 | / | MN032180 | MN032247 |
| *Episoriculus macrurus* | SAF13198 | NJ13032 | Lushui, Yunnan | 98.70969 | 25.98614 | 2719 | MK962170 | MN032126 | MN032184 | MN032251 |
| *Episoriculus macrurus* | SAF13256 | NJ13090 | Gongshan, Yunnan | 98.44691 | 27.77396 | 3320 | MK962165 | / | MN032179 | MN032246 |
| *Episoriculus macrurus* | SAF13288 | NJ13122 | Gongshan, Yunnan | 98.50351 | 27.79702 | 3050 | MK962167 | / | MN032181 | MN032248 |
| *Episoriculus macrurus* | SAF13197 | NJ13031 | Lushui, Yunnan | 98.70969 | 25.98614 | 2719 | MK962163 | / | MN032177 | / |
| *Episoriculus macrurus* | SAF07587 | CY17 | Chayu, Tibet | 97.01826 | 28.77404 | 2900 | MK962236 | MN032173 | / | / |
| *Episoriculus macrurus* | SAF09629 | JJSA506 | Jinjiashan, Sichuan | 102.75222 | 30.84041 | 2620 | MK962195 | / | MN032205 | MN032274 |
| *Episoriculus macrurus* | SAF181173 | WL18246 | Wanglang, Sichuan | 104.1342 | 32.94794 | 2570 | MK962179 | MN032134 | / | MN032260 |
| *Episoriculus macrurus* | SAF06900 | ELSTNPB02002 | Erlangshan, Sichuan | 102.29916 | 29.88547 | 2450 | MK962175 | MN032131 | MN032188 | MN032256 |
| *Episoriculus macrurus* | SAF06829 | ELSMYPA02004 | Erlangshan, Sichuan | 102.28433 | 29.86774 | 2780 | MK962162 | / | MN032176 | MN032244 |
| *Episoriculus macrurus* | SAF11599 | BLG031 | Balanggou, Sichuan | 101.91481 | 30.40052 | 2820 | MK962187 | MN032139 | MN032197 | / |
| *Episoriculus macrurus* | SAF06932 | ELS260B02001 | Erlangshan, Sichuan | 102.31112 | 29.87314 | 2230 | MK962173 | / | MN032187 | MN032254 |
| *Episoriculus macrurus* | SAF06874 | ELSTNPA02002 | Erlangshan, Sichuan | 102.29953 | 29.88633 | 2430 | MK962176 | / | MN032189 | MN032257 |
| *Episoriculus macrurus* | SAF181086 | WL18159 | Wanglang, Sichuan | 104.1437 | 32.928634 | 2500 | MK962180 | MN032135 | / | MN032261 |
| *Episoriculus macrurus* | SAF181215 | WL18288 | Wanglang, Sichuan | 117.1342 | 45.94794 | 2583 | MK962183 | / | MN032193 | MN032263 |
| *Episoriculus macrurus* | SAF181127 | WL18200 | Wanglang, Sichuan | 104.1459 | 32.927797 | 2500 | MK962189 | MN032140 | MN032199 | MN032268 |
| *Episoriculus macrurus* | SAF181126 | WL18199 | Wanglang, Sichuan | 104.1459 | 32.927797 | 2500 | MK962188 | / | MN032198 | MN032267 |
| *Episoriculus macrurus* | SAF181065 | WL18138 | Wanglang, Sichuan | 104.1437 | 32.928634 | 2500 | MK962185 | / | MN032195 | MN032265 |
| *Episoriculus macrurus* | SAF181130 | WL18203 | Wanglang, Sichuan | 104.1459 | 32.927797 | 2500 | MK962184 | MK962184 | MN032194 | MN032264 |
| *Episoriculus macrurus* | SAF181129 | WL18202 | Wanglang, Sichuan | 104.1459 | 32.927797 | 2500 | MK962190 | MN032141 | MN032200 | MN032269 |
| *Episoriculus macrurus* | SAF09655 | JJSA532 | Balanggou, Sichuan | 102.75311 | 30.84053 | 2620 | MK962193 | MN032143 | MN032203 | MN032272 |
| *Episoriculus macrurus* | SAF181137 | WL18137 | Wanglang, Sichuan | 104.1459 | 32.927797 | 2500 | MK962182 | MN032136 | MN032192 | / |
| *Episoriculus macrurus* | SAF06725 | ELSLCA03001 | Erlangshan, Sichuan | 102.26703 | 29.85678 | 2800 | MK962172 | / | MN032186 | MN032253 |
| *Episoriculus macrurus* | SAF181394 | WL18467 | Wanglang, Sichuan | 103.9963 | 32.96181 | 3000 | MK962174 | MN032130 | / | MN032255 |
| *Episoriculus macrurus* | SAF181128 | WL18201 | Wanglang, Sichuan | 104.1459 | 32.927797 | 2500 | MK962181 | / | MN032191 | MN032262 |
| *Episoriculus macrurus* | SAF15165 | LS15023 | Lushan, Sichuan | 103.01808 | 30.68135 | 2424 | MK962194 | / | MN032204 | MN032273 |
| *Episoriculus macrurus* | SAF06730 | ELSDGA01003 | Erlangshan, Sichuan | 102.2584 | 29.85416 | 2700 | MK962196 | / | MN032206 | MN032275 |
| *Episoriculus macrurus* | SAF06767 | ELSBTDBS01013 | Erlangshan, Sichuan | 102.26283 | 29.85202 | 2500 | MK962161 | / | MN032175 | MN032243 |
| *Episoriculus macrurus* | JJSA137 | JJSA137 | Jiajinshan, Sichuan |  |  |  | MK962192 | MN032142 | MN032202 | MN032271 |
| *Episoriculus macrurus* | SAF181202 | WL18275 | Wanglang, Sichuan | 104.1342 | 32.94794 | 2570 | MK962178 | MN032133 | / | MN032259 |
| *Episoriculus macrurus* | SAF09303 | JJSA443 | Jiajinshan, Sichuan | 10268985 | 30.84276 | 3440 | MK962186 | MN032138 | MN032196 | MN032266 |
| *Episoriculus macrurus* | HLG01017 | HLG01017 | Hailuogou, Sichuan |  |  |  | MK962191 | / | MN032201 | MN032270 |
| *Episoriculus macrurus* | SAF06177 | MGDW0502 | Meigu, Sichuan | 103.27607 | 28.7063 | 2900 | MK962177 | MN032132 | MN032190 | MN032258 |
| *Episoriculus soluensis* | SAF13503 | XZ13041 | Nyalam, Tibet | 85.99854 | 28.0815 | 3200 | MK962201 | MN032146 | MN032210 | MN032278 |
| *Episoriculus soluensis* | SAF14536 | XZ14002 | Yadong, Tibet | 88.994744 | 27.540042 | 4304 | MK962157 | / | / | / |
| *Episoriculus soluensis* | SAF14537 | XZ14003 | Yadong, Tibet | 88.994744 | 27.540042 | 4304 | MK962158 | / | / | / |
| *Episoriculus soluensis* | SAF14538 | XZ14004 | Yadong, Tibet | 88.994744 | 27.540042 | 4304 | MK962159 | / | / | / |

\* Indicates that this sequence was downloaded from NCBI.

Following He *et al*. [15] and Chen *et al*. [16,17], we amplified the complete mitochondrial cytochrome b (*CYTB*) and three partial nuclear genes (apolipoprotein B (*APOB*), recombination-activating gene 2 (*RAG2*), and breast cancer 1(*BRCA1*)). Primer sets are detailed in Table 3[15, 18–20]. PCR amplifications were carried out in a 25 µl reaction volume mixture containing 12.5 µl of 2×Taq Master Mix (Vazyme, Nanjing, China), 1 µl of each primer, 1 µl of genomic DNA, and 9.5 µl of double-distilled water. PCR conditions for *CYTB* amplifications consisted of an initial denaturing step at 94°C for 5 min followed by 38 cycles of denaturation at 94°C for 45 s, annealing at 49°C for 45 s, an extension at 72°C for 90 s, and a final extension step at 72°C for 12 min. PCR conditions for nuclear genes were basically the same as those for *CYTB*, with a few modifications (annealing temperatures for each nuclear gene were *APOB* (49°C), *BRCA1* (51°C), and *RAG2* (52°C)). PCR products were checked on a 1.0% agarose gel and purified by ethanol precipitation. Purified PCR products were directly sequenced using the BigDye Terminator Cycle Kit v 3.1 (Applied Biosystems, Foster City, CA, USA) and an ABI 310 Analyzer (Applied Biosystems).

**Table 3.**
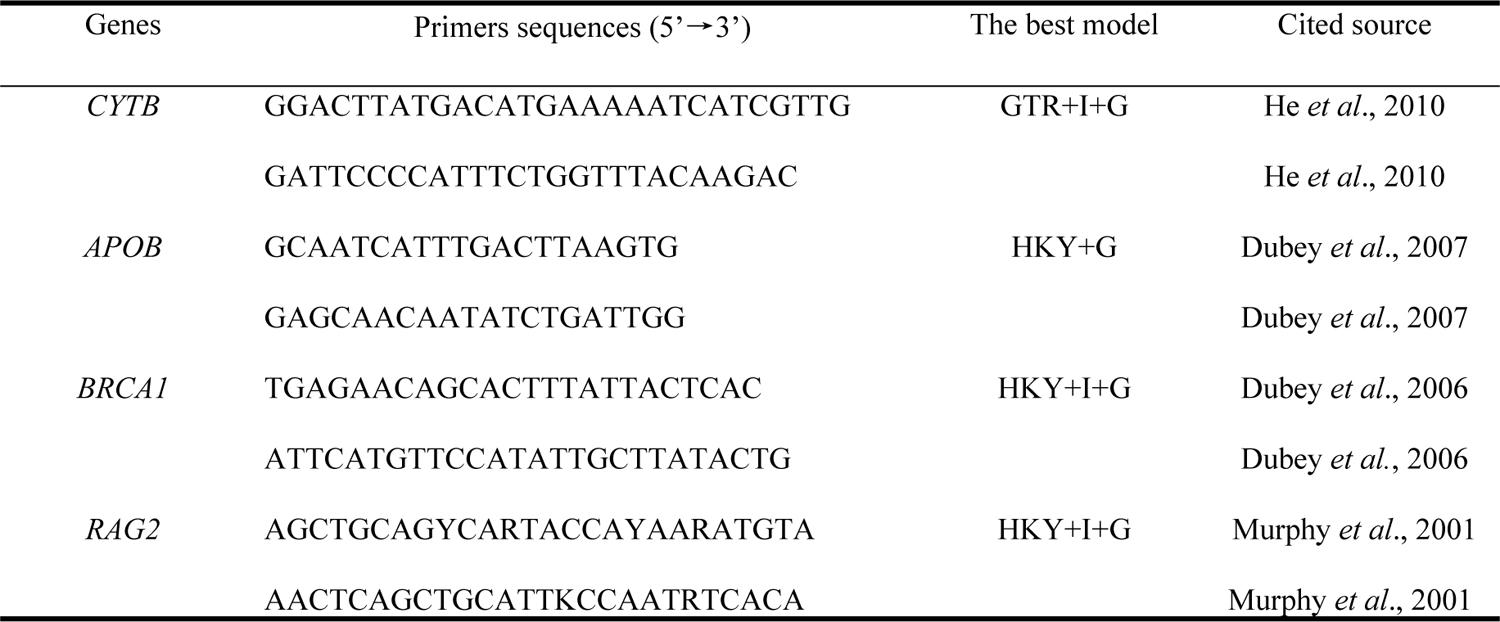
Gene symbol, primer sequences and the best model of evolution for each gene segments used in the study.

To test phylogenetic relationships within the genus *Episoriculus*, sequences of these four genes from other Nectogalini species generated in previous studies [2, 15, 16] were downloaded from GenBank (Table 4).

**Table 4.**
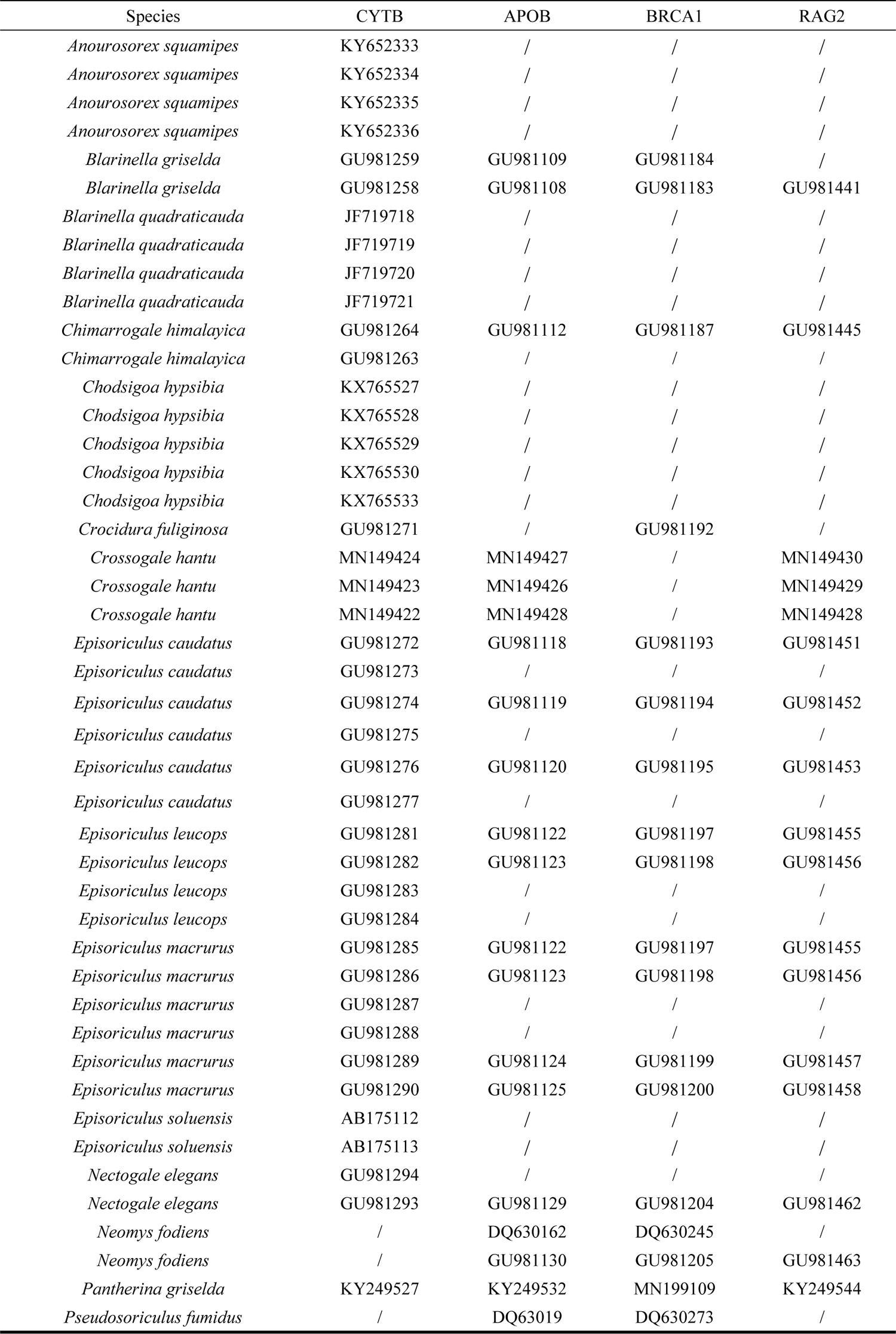
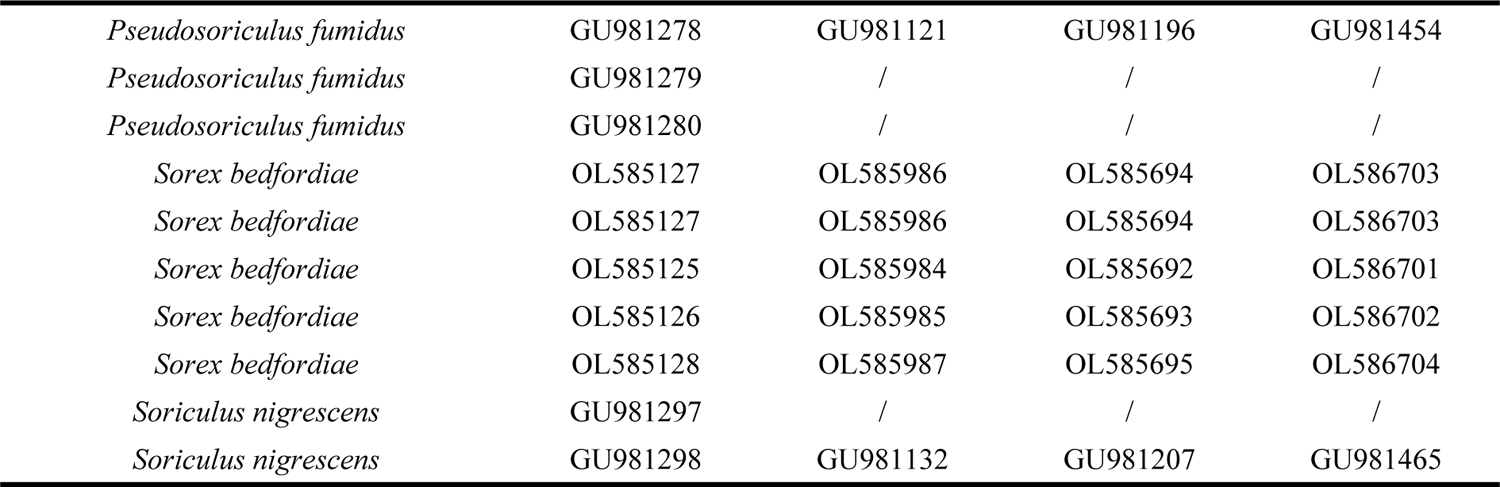
GenBank accession numbers of download sequence from NCBI.

| Species | CYTB | APOB | BRCA1 | RAG2 |
| --- | --- | --- | --- | --- |
| *Anourosorex squamipes* | KY652333 | / | / | / |
| *Anourosorex squamipes* | KY652334 | / | / | / |
| *Anourosorex squamipes* | KY652335 | / | / | / |
| *Anourosorex squamipes* | KY652336 | / | / | / |
| *Blarinella griselda* | GU981259 | GU981109 | GU981184 | / |
| *Blarinella griselda* | GU981258 | GU981108 | GU981183 | GU981441 |
| *Blarinella quadraticauda* | JF719718 | / | / | / |
| *Blarinella quadraticauda* | JF719719 | / | / | / |
| *Blarinella quadraticauda* | JF719720 | / | / | / |
| *Blarinella quadraticauda* | JF719721 | / | / | / |
| *Chimarrogale himalayica* | GU981264 | GU981112 | GU981187 | GU981445 |
| *Chimarrogale himalayica* | GU981263 | / | / | / |
| *Chodsigoa hypsibia* | KX765527 | / | / | / |
| *Chodsigoa hypsibia* | KX765528 | / | / | / |
| *Chodsigoa hypsibia* | KX765529 | / | / | / |
| *Chodsigoa hypsibia* | KX765530 | / | / | / |
| *Chodsigoa hypsibia* | KX765533 | / | / | / |
| *Crocidura fuliginosa* | GU981271 | / | GU981192 | / |
| *Crossogale hantu* | MN149424 | MN149427 | / | MN149430 |
| *Crossogale hantu* | MN149423 | MN149426 | / | MN149429 |
| *Crossogale hantu* | MN149422 | MN149428 | / | MN149428 |
| *Episoriculus caudatus* | GU981272 | GU981118 | GU981193 | GU981451 |
| *Episoriculus caudatus* | GU981273 | / | / | / |
| *Episoriculus caudatus* | GU981274 | GU981119 | GU981194 | GU981452 |
| *Episoriculus caudatus* | GU981275 | / | / | / |
| *Episoriculus caudatus* | GU981276 | GU981120 | GU981195 | GU981453 |
| *Episoriculus caudatus* | GU981277 | / | / | / |
| *Episoriculus leucops* | GU981281 | GU981122 | GU981197 | GU981455 |
| *Episoriculus leucops* | GU981282 | GU981123 | GU981198 | GU981456 |
| *Episoriculus leucops* | GU981283 | / | / | / |
| *Episoriculus leucops* | GU981284 | / | / | / |
| *Episoriculus macrurus* | GU981285 | GU981122 | GU981197 | GU981455 |
| *Episoriculus macrurus* | GU981286 | GU981123 | GU981198 | GU981456 |
| *Episoriculus macrurus* | GU981287 | / | / | / |
| *Episoriculus macrurus* | GU981288 | / | / | / |
| *Episoriculus macrurus* | GU981289 | GU981124 | GU981199 | GU981457 |
| *Episoriculus macrurus* | GU981290 | GU981125 | GU981200 | GU981458 |
| *Episoriculus soluensis* | AB175112 | / | / | / |
| *Episoriculus soluensis* | AB175113 | / | / | / |
| *Nectogale elegans* | GU981294 | / | / | / |
| *Nectogale elegans* | GU981293 | GU981129 | GU981204 | GU981462 |
| *Neomys fodiens* | / | DQ630162 | DQ630245 | / |
| *Neomys fodiens* | / | GU981130 | GU981205 | GU981463 |
| *Pantherina griselda* | KY249527 | KY249532 | MN199109 | KY249544 |
| *Pseudosoriculus fumidus* | / | DQ63019 | DQ630273 | / |
| *Pseudosoriculus fumidus* | GU981278 | GU981121 | GU981196 | GU981454 |
| *Pseudosoriculus fumidus* | GU981279 | / | / | / |
| *Pseudosoriculus fumidus* | GU981280 | / | / | / |
| *Sorex bedfordiae* | OL585127 | OL585986 | OL585694 | OL586703 |
| *Sorex bedfordiae* | OL585127 | OL585986 | OL585694 | OL586703 |
| *Sorex bedfordiae* | OL585125 | OL585984 | OL585692 | OL586701 |
| *Sorex bedfordiae* | OL585126 | OL585985 | OL585693 | OL586702 |
| *Sorex bedfordiae* | OL585128 | OL585987 | OL585695 | OL586704 |
| *Soriculus nigrescens* | GU981297 | / | / | / |
| *Soriculus nigrescens* | GU981298 | GU981132 | GU981207 | GU981465 |

### Sequence analyses

All *CYTB* sequences were aligned and examined. Screening for heterozygote nuclear gene fragments was performed in Mega 5 [21]. For analysis, we concatenated the three nuclear genes following [15]. Using all sequence data, phylogenetic analyses were conducted on: 1) *CYTB* data, 2) concatenated sequences for the three nuclear genes, and 3) each nuclear gene. Modeltest v 3.7 [22] was used to select the best-fitting evolutionary model, based on the Akaike Information Criterion in Table 3. MrBayes v 3.1.2 [23] was used for Bayesian analysis. *Crocidura fuliginosa* was selected as the outgroup. Each run was carried out with four Monte Carlo Markov chains (MCMCs), and 10,000,000 generations for single gene datasets and 30,000,000 generations for concatenated gene datasets. All runs were sampled every 10,000 generations. Convergences of runs were accepted when the average standard deviation of split frequencies was < 0.01. Ultrafast bootstrap values (UFBoot) of ≥ 95 and posterior probabilities (PP) of ≥ 0.95 were considered strong support [23].

### Species tree and species delimitation

To appraise current taxonomic systems based on morphology, and to evaluate trees derived from phylogenetic analyses, we used species delimitation and species tree construction based on abductive theory. We estimated the *BEAST coalescent species tree in Beast v 1.7.5 using phased nuclear and mitochondrial genes [24–26]. Model settings were selected with reference to the optimal replacement model of each gene (Table 3). Each analysis was run for 100 ×10^6^ generations and sampled every 10,000 generations. Posterior distribution and effective sample size of each parameter were calculated using Tracer v 1.6. *BEAST analyses were repeated four times to ensure convergence on the same posterior distribution.

“Splits” (Species Limits by Threshold Statistics) v 1.0 14 was used for species delimitation in the context of R statistics. The program defines species using the generalized mixed Yule-coalescent model (GMYC) [27, 28]. Analysis requires a gene tree that has been corrected by a molecular clock as a reference; we use the bifurcated time tree constructed with Beast v 1.7.5 as basic data for this analysis.

BPP v 2.2 was used to progress species delimitation [29]. Analyses of species boundaries were limited to *E. caudatus*, *E. sacratus*, and *E. umbrinus*; analyses used the combined nuclear gene data set. Since this program also requires a prior guide tree, and the topological structure of the guide tree will impact the result of species delimitation, we used the a forementioned species tree built by BEAST v 1.7.5 as the guide tree for BPP analysis [30]. For BPP, we set the Gamma prior distribution of the population size parameter (θs) to G (6, 6,000), and the initial differentiation time parameter (τ0) of the species tree to G (4, 1,000). Then 12 parameter combinations were generated using algorithms 0 and 1, and combining the values of Locusrate = 1 or Heredity = 1. The above 12 operations were performed on the two data groups; each operation was set to 1,000,000 generations of reverse-jump MCMC, and samples were taken every 10 generations; the first 10,000 generations were discarded [31].

Automatic barcode gap discovery (ABGD) software was used to divide samples based on genetic distance; samples within the same group were identified as one species [32]. *CYTB* sequences of the sample online submission to ABGD website (http://www.abi.snv.Jussleu.fr/publicabgd/abgdweb.hml), the prior intraspecific divergence (P) ranged 0.001–0.1, and the minimum relative gap width (X) was 0.5. We also used the Kimura 2-parameter (K2p) distance with 10,000 bootstrap replicates to summarize sequence divergences based on *CYTB* in MEGA5 [33].

### Morphometrics

Because we have found no evidence for sexual duality in shrews in related taxa, we do not consider sex when selecting specimens and skulls. All samples used for analysis were adults with intact skulls. Specimens of *Episoriculus* used for study have been deposited in the Sichuan Academy of Forestry, and Kunming Institute of Zoology. A total of 56 complete skulls of adult intact specimens were assigned to *E. macrurus* (17), *E. caudatus* (11), *E. umbrinus* (8), *E. leucops* (7), *E. sacratus* (6), and *E. soluensis* (3). Details of localities and museums are listed in Table S1.

Measurements including head and body length (HBL), tail length (TL), hind foot length (HFL), and ear length (EL) were recorded from specimen labels or field notes. Eleven craniomandibular variables were measured by digital caliper (to 0.01 mm) from 56 specimens: condylox incisive length (CIL), cranial height (CH), cranial breadth (CB), interorbital breadth (IBO), palatoincisive length (PIL), postpalatal length (PPL), maxillary breadth (MB), upper toothrow length (UTR), maximum width across the upper second molars (M^2^–M^2^), mandibular length (ML), and lower toothrow length (LTR). Eleven measurements were used for principal component analysis (PCA) in SPSS 22.0 (SPSS Inc., USA). Sample localities and measurements for each specimen are presented in Table S1. Measuring methods followed Chen *et al*. [16], Yang *et al*. [34].

Cranial measurements were analyzed by PCA in SPSS v19.0 (SPSS Inc., USA). PCAs were conducted on log_10_ transformed variables on two data sets. Before analysis, the Kaiser– Meyer–Olkin (KMO) test (to check correlations or partial correlations between variables), and Bartlett’s sphere test (to determine if the correlation matrix is an identity matrix) were performed.

## Results

### Phylogenetic analysis

Bayesian reconstruction using *CYTB* revealed eight monophyletic clades of *Episoriculus*, corresponding to *E. macrurus*, *E. soluensis*, *E. leucops*, *E. caudatus*, *E. umbrinus*, *E. sacratus*, and *E.* sp. (Fig. 2a). *Pseudosoriculus fumidus* clustered with *Chodsigoa* with strong support (PP = 0.95). *Episoriculus macrurus* represented a basal lineage of Nectogalini with strong support (PP = 1.00), and *E. soluensis*, *E. leucops*, *E. sacratus* and *E.* sp. formed a separate, strongly supported lineage (PP = 0.95–1.00). *Episoriculus umbrinus* and *E. caudatus*, located at the tip of this tree, formed sister clades, that were less-well supported (PP = 0.75).

**Fig. 2.**
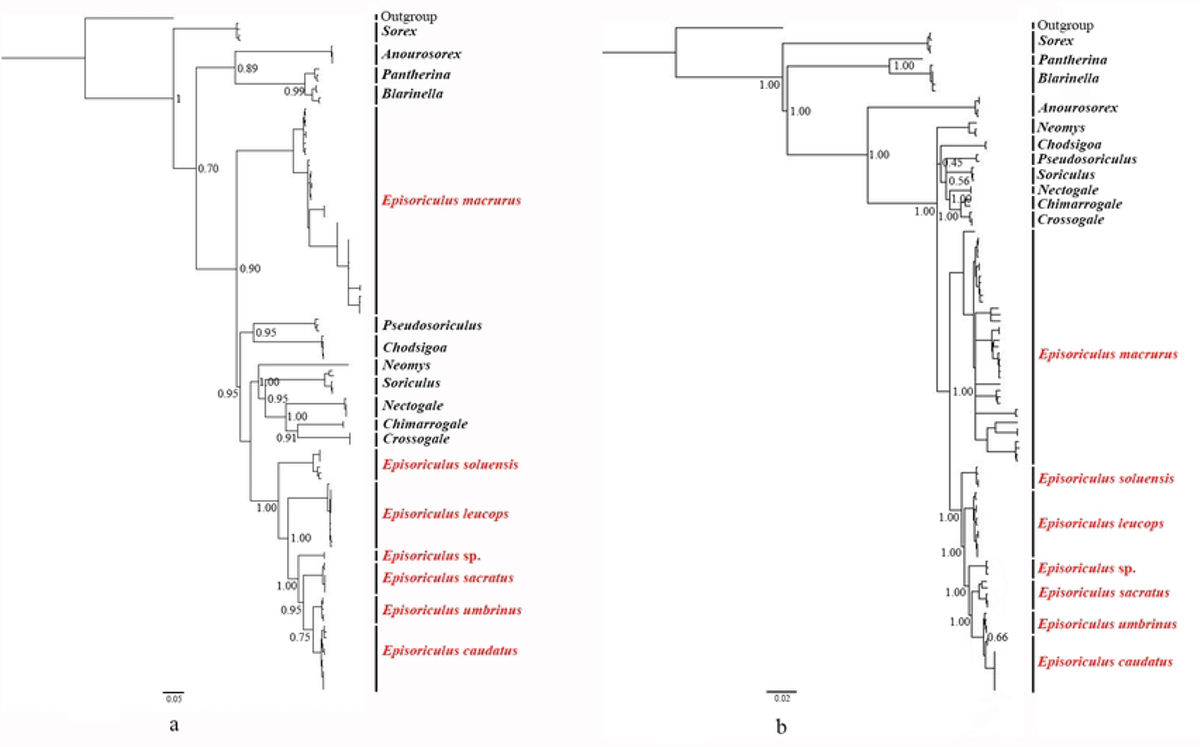
Bayesian phylogenetic analyses based on (a) *CYTB*, and (b) three concatenated nuclear genes. Numbers at nodes refer to Bayesian posterior probabilities. Scale bars represent substitutions per site. This is the Fig. 2 legend.

Bayesian reconstruction using the three concatenated nuclear genes also revealed eight monophyletic clades (Fig. 2b), but with a slightly different topological structure to the *CYTB* tree. *Pseudosoriculus fumidus* mixed with *Chodsigoa* and *Soriculus*, with weak support. All species of *Episoriculus* formed a monophyletic clade, with *E. macrurus* at its base. Lineages of *E. soluensis*, *E. leucops*, *E. sacratus*, and *E.* sp. were strongly supported (PP = 1.00). *Episoriculus umbrinus* and *E. caudatus* formed sister clades at the tree’s tip, although support for them was weak (PP = 0.66).

Structures of three individual nuclear gene trees differed from the CYTB and three concatenated nuclear gene trees (Fig. 3), with some nodes having very low support. *Pseudosoriculus fumidus* did not cluster with Episoriculus. E*pisoriculus macrurus*, *E. soluensis*, *E. leucops*, *E.* sp., and *E. sacratus* remained monophyletic with strong support based on *APOB* and *RAG2* (PP = 1.00) genes, and *E. umbrinus* and *E. caudatus* formed a sister group with weak support. *Episoriculus sacratus*, *E. umbrinus* and *E. caudatus* were mixed on the tree based on the *BRCA1* gene, with very low support.

**Fig 3.**
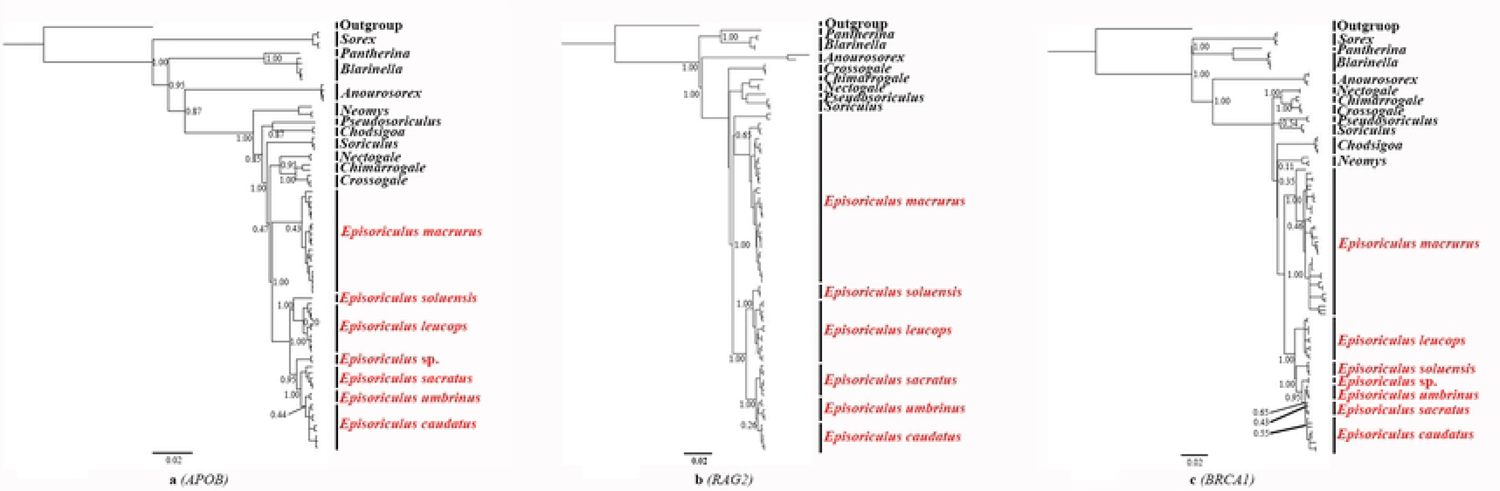
Bayesian phylogenetic analyses based on (a) *APOB*, (b) *RAG2*, and (c) *BRCA1* genes. Numbers at nodes refer to Bayesian posterior probabilities. Scale bars represent substitutions per site. This is the Fig. 3 legend.

### Species delimitation

The topology of *BEAST species’ trees differed slightly from those of mitochondrial and nuclear genes (Fig. 4). *Episoriculus macrurus*, *E*. *soluensis*, *E. leucops*, *E. sacratus*, and *E.* sp. also had high support in these trees (PP = 1.00). *Episoriculus umbrinus* and *E. caudatus* were sister clades, but with weak support in the tree (PP = 0.75). This result did not support recognizing *E. umbrinus* as a distinct species.

**Fig 4.**
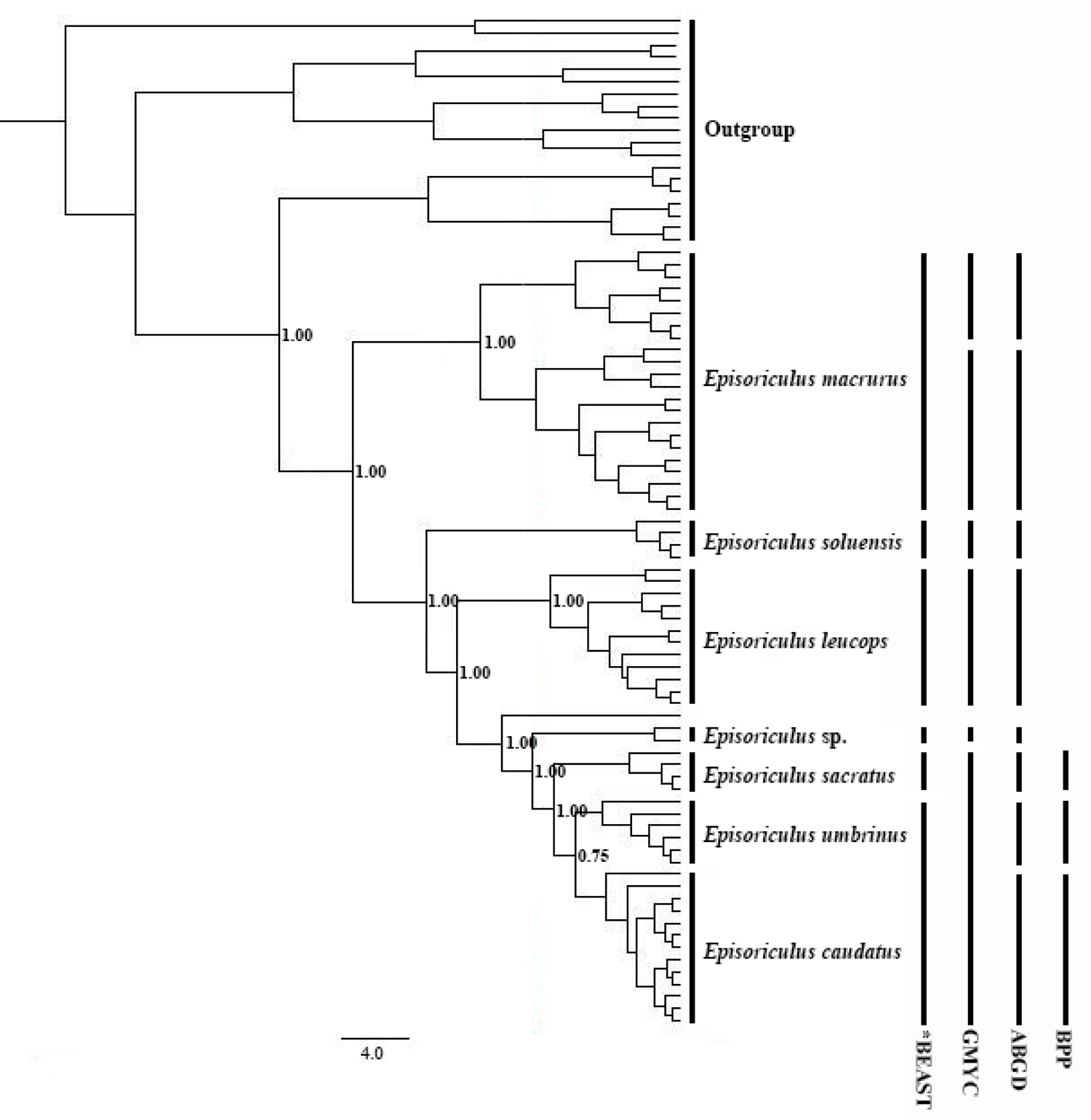
Results of species delimitation using splits, GMYC, BPP, and species trees reconstructed using the *BEAST model. Node numbers indicate Bayesian posterior probabilities supporting each clade as two putative species. This is the **Fig. 4** legend

GMYC analysis reveals five clades as valid species (Fig. 4), of which *E. soluensis*, *E. leucops,* and *E.* sp. are separate, and *E. macrurus* comprised two species, with individuals from Sichuan and Yunnan provinces differing. This analysis suggests that *E. caudatus*, *E. sacratus* and *E. umbrinus* are conspecific.

For the ABGD analysis, the transition/transversion value (3.5) first calculated by Mega 5 was used as the starting parameter, with 0.5, 1.0, 1.5 and 2.0 used as relative gap widths. The 81 samples divided into eight species: *E. caudatus*, *E. macrurus*, *E. sacratus*, *E. soluensis*, *E. leucops*, *E. macrurus* (Sichuan samples), *E. macrurus* (Yunnan samples), and *E.* sp. (Fig. 4, Table S2).

BPP analysis revealed *E. caudatus* and *E. umbrinus* to be separate species in 12 groups of BPP data based on the combined nuclear gene data set, with support for *E. sacratus* being a valid taxon also being high (Fig. 4, Table S3).

Kimura-2-parameter (K2p) distances between *Episoriculus* species ranged 0.027–0.160 (Table 5). The average K2p distance between *P. fumidus* and *Episoriculus* species was 0.177. The K2p distance between *E. caudatus* and *E. sacratus* was 0.067, and between *E. caudatus* and *E. umbrinus*, 0.027, and *E. sacratus* and *E. umbrinus*, 0.071.

**Table 5.** The Kimura-2-parameter distances between *Episoriculus* species based on the *CYTB* gene.

| No. | Species | 1 | 2 | 3 | 4 | 5 | 6 | 7 |
| --- | --- | --- | --- | --- | --- | --- | --- | --- |
| 1 | *Episoriculus caudatus* |  |  |  |  |  |  |  |
| 2 | *Episoriculus sacratu*s | 0.066 |  |  |  |  |  |  |
| 3 | *Episoriculus umbrinus* | 0.027 | 0.071 |  |  |  |  |  |
| 4 | *Episoriculus* sp. | 0.083 | 0.088 | 0.099 |  |  |  |  |
| 5 | *Episoriculus soluensis* | 0.113 | 0.121 | 0.128 | 0.129 |  |  |  |
| 6 | *Episoriculus leucops* | 0.112 | 0.133 | 0.131 | 0.126 | 0.139 |  |  |
| 7 | *Episoriculus macrurus* | 0.149 | 0.156 | 0.160 | 0.148 | 0.159 | 0.154 |  |
| 8 | *Pseudosoriculus fumidus* | 0.181 | 0.184 | 0.192 | 0.177 | 0.163 | 0.189 | 0.154 |

### Morphology

Morphological data (HBL, TL, HFL, EL, and body weight (BW)) of 56 specimens with intact skulls are presented in Table 6. The TL of *E. macrurus* ranged 71–106mm, while values for congeners ranged 46–82.5 mm. The HFL of *E. macrurus* was 15–16 mm, while for other species it was 12–13 mm. *Episoriculus macrurus* had the lowest HBL/TL ratio (0.64), and its TL was ∼1.5 times its HBL, while the HBL of congeners was approximately equal to or greater than TL. *Episoriculus leucops* had the largest body, and *E. sacratus* differed from *E. caudatus* and *E. umbrinus* in HBL/TL values. Morphological indices are detailed in Table 7, and complete measurement data are provided in Table S1.

**Table 6.** The results of body morphologic measurements.

| Species | N | BW | HBL | TL | HFL | HBL / TL |
| --- | --- | --- | --- | --- | --- | --- |
| *E. caudatus* | 14 | 5.000±0.327 | 56.750±1.858 | 54.750±0.491 | 12.125±0.125 | 1.04 |
| *E. sacratu* | 6 | 5.666±0.211 | 57.134±0.632 | 62.166±2.249 | 13.500±0.671 | 0.92 |
| *E. umbrinus* | 9 | 5.000±0.239 | 61.333±0.816 | 60.222±0.547 | 13.000±0.231 | 1.01 |
| *E. leucops* | 7 | 7.714±0.286 | 62.714±0.056 | 60.143±1.534 | 13.143±0.143 | 1.04 |
| *E. macrurus* | 17 | 5.000±0.413 | 58.158±1.732 | 91.333±1.201 | 15.667±0.333 | 0.64 |
| *E. soluensis* | 4 | 6.000±0.210 | 64.333±0.666 | 61.333±0.088 | 12.000±0.531 | 1.04 |

**Table 7.** Morphological measurement data of *Episoriculus* species skulls.

| Species | N | CIL | IOB | CB | BH | MB | PIL | PPL | UTR | M<sup>2</sup>-M<sup>2</sup> | ML | LTR |
| --- | --- | --- | --- | --- | --- | --- | --- | --- | --- | --- | --- | --- |
| *E. caudatus* | 14 | 18.327 | 3.311 | 8.769 | 5.907 | 1.391 | 8.104 | 8.067 | 8.1464 | 4.525 | 11.258 | 7.2657 |
|  |  | ±0.057 | ±0.044 | ±0.374 | ±0.068 | ±0.018 | ±0.059 | ±0.039 | ±0.058 | ±0.038 | ±0.056 | ±0.058 |
| *E. sacratu* | 6 | 18.206 | 3.430 | 8.578 | 6.030 | 1.381 | 7.840 | 8.035 | 7.773 | 4.565 | 11.036 | 6.981 |
|  |  | ±0.593 | ±0.052 | ±0.082 | ±0.100 | ±0.237 | ±0.056 | ±0.103 | ±0.073 | ±0.029 | ±0.090 | ±0.046 |
| *E. umbrinus* | 8 | 18.286 | 3.523 | 8.513 | 5.628 | 1.371 | 8.208 | 8.126 | 8.052 | 4.457 | 11.1789 | 7.3933 |
|  |  | ±0.070 | ±0.054 | ±0.045 | ±0.045 | ±0.024 | ±0.063 | ±0.088 | ±0.046 | ±0.049 | ±0.077 | ±0.0763 |
| *E. leucops* | 7 | 19.307 | 3.708 | 9.068 | 5.858 | 1.391 | 8.658 | 8.876 | 8.416 | 4.564 | 11.693 | 7.354 |
|  |  | ±0.111 | ±0.377 | ±0.448 | ±0.046 | ±0.341 | ±0.036 | ±0.797 | ±0.082 | ±0.062 | ±0.130 | ±0.147 |
| *E. macrurus* | 18 | 17.577 | 3.309 | 8.456 | 5.761 | 1.290 | 7.633 | 7.813 | 7.565 | 4.184 | 10.618 | 6.781 |
|  |  | ±0.060 | ±0.191 | ±0.057 | ±0.050 | ±0.012 | ±0.051 | ±0.079 | ±0.056 | ±0.045 | ±0.051 | ±0.036 |
| *E. soluensis* | 4 | 17.896 | 3.603 | 9.327 | 5.763 | 1.706 | 8.067 | 7.976 | 7.767 | 4.873 | 12.320 | 7.497 |
|  |  | ±0.003 | ±0.052 | ±0.097 | ±0.233 | ±0.328 | ±0.296 | ±0.024 | ±0.084 | ±0.003 | ±0.055 | ±0.084 |

Bartlett’s test rejected the null hypothesis (χ^2^ = 568.01, P = 0.000), indicating that the data were spherical and variables were somewhat independent of each other. A KMO of 0.813 indicated a strong correlation existed among the various skull data, which was suitable for factor analysis. Two principal components explaining 74.69% of morphological variation were extracted from the analysis. Factor loading values were most positive, indicating that it was mainly related to overall skull size (Table 8). Features with factor loads > 0.8 included PIL, ML, CIL, UTR, and LTR.

**Table 8.** Character loadings, eigenvalues, and proportion of variance explained by the first two axes (PC 1 and PC 2) of a principal component analysis using the log10-transformed measurements of *Episoriculus*. The meanings of variable abbreviations are given in the Materials and Methods Section.

| Measurement | Principal component (PC) |  |
| --- | --- | --- |
|  | 1 | 2 |
| PIL | 0.888 | -0.340 |
| ML | 0.886 | 0.216 |
| CIL | 0.876 | -0.247 |
| UTR | 0.841 | -0.279 |
| LTR | 0.804 | -0.003 |
| CB | 0.750 | 0.290 |
| M<sup>2</sup>-M<sup>2</sup> | 0.733 | 0.502 |
| PPL | 0.704 | -0.409 |
| IOB | 0.682 | -0.258 |
| MB | 0.592 | 0.722 |
| CH | 0.030 | 0.537 |
| Variance explained | 61.022 | 13.664 |

Using PC 1 and PC 2 maps (Fig. 5), *E. macrurus* plotted in the positive region of PC 2, while other species were mainly in the negative region of PC 2. The larger *E. leucops* plotted in the negative region of PC 2, and *E. soluensis* plotted in the positive region of PC 2. *Episoriculus sacratus* was distinguished with *E. umbrinus* and *E. caudatus*, with the latter two species being mixed and not effectively differentiated.

**Fig. 5.**
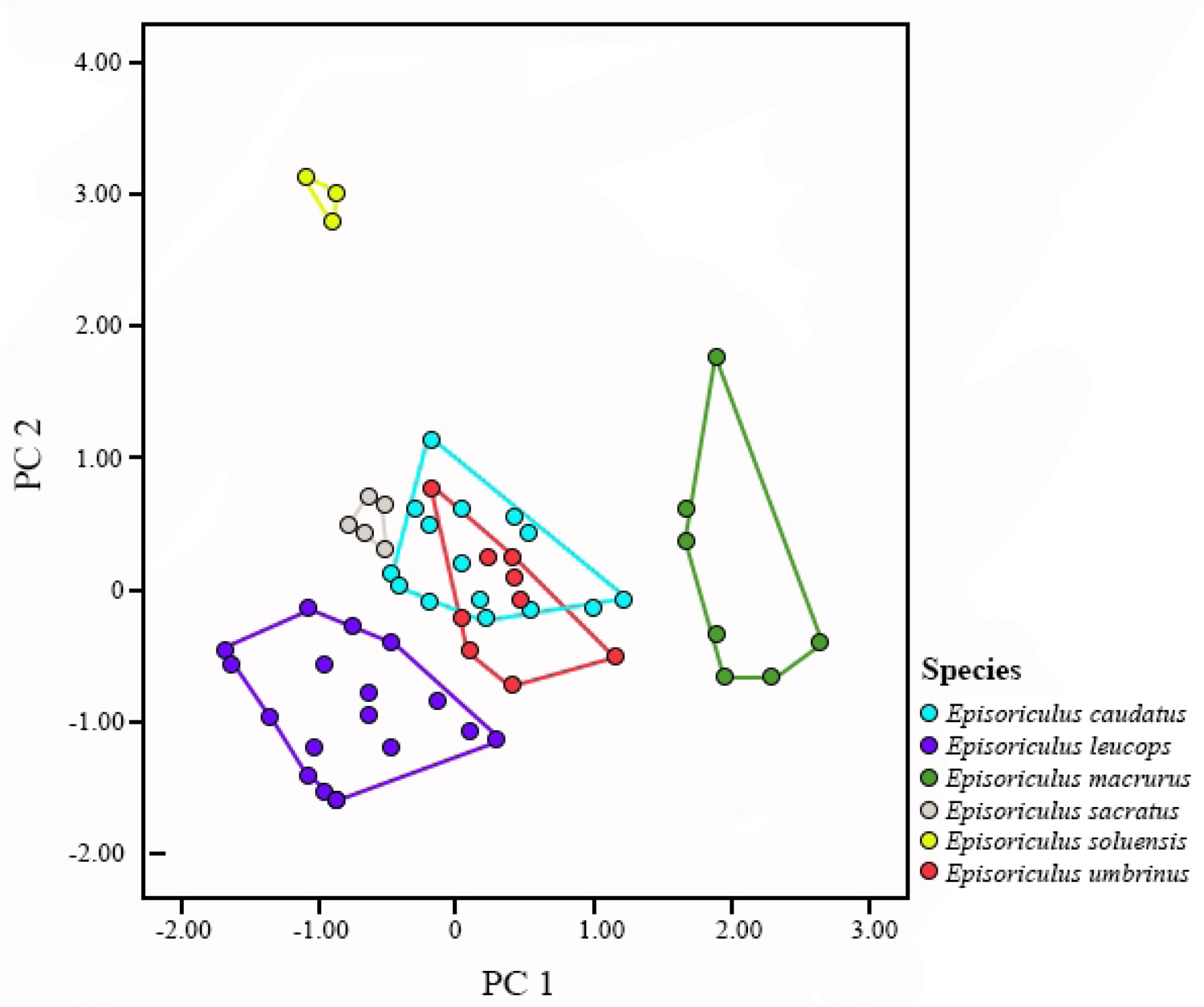
Results of principal component analysis of *Episoriculus* taxa based on 19 log_10_ transformed craniodental measurements. This is the Fig. 5 legend

Compared to skulls of other *Episoriculus* species (Fig. 6), the braincase of *E. macrurus* is more dome-shaped, the rostrum is shorter; and the upper unicuspids are quadrangular and wider than long (those of other species are similarly sized). Compared to skulls of *E. caudatus*, the frontal region of the skull of *E. sacratus* is more arched, and the posterior cusp of its upper incisor is lower than its first unicuspid, whereas the height of the posterior cusp of the upper incisor and first unicuspid of *E. caudatus* are similar.

**Fig. 6.**
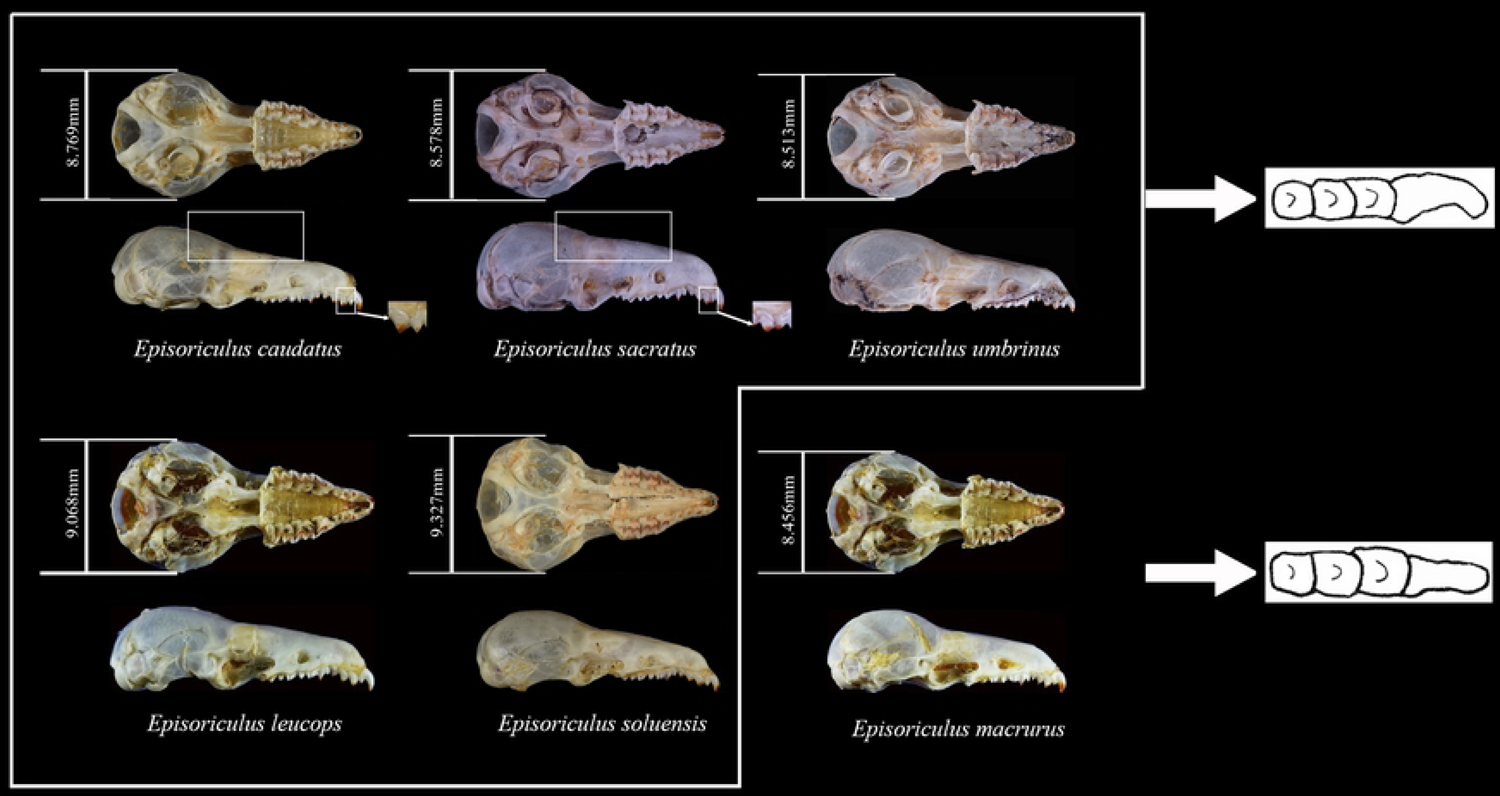
Comparison of *Episoriculus* skulls. This is the Fig. 6 legend.

## Discussion

### Differences between mitochondrial and nuclear genes

We report inconsistencies in phylogenies of *Episoriculus* and related species that have been reconstructed from mitochondrial and nuclear gene data. Similar conflicting phylogenetic signals have been reported in other studies [35], and could be explained by differences in genetic background [36], ancient hybridization [37], incomplete lineage sorting [38], adaptive evolution, and burst Formula speciation [39]. While neither mtDNA nor nDNA alone resolved phylogenetic relationships in the genus *Episoriculus*, combining data from these two genetic pathways did improve results. Species tree construction in the anadromous framework also produced a consistent topology with high statistical support. Therefore, we deem that a combined approach using mitochondrial and nuclear gene information is more appropriate for resolving phylogenetic relationships in the genus *Episoriculus*.

### Species richness

Thomas [40] described six specimens from Mount Emei as *E. sacratus*, and considered it most likely the local representative of *E. caudatus*, from which they differed in having a much smaller braincase. Allen [41] described the subspecies *E. caudatus umbrinus* from Mucheng, Yunnan, which was most alike *E. sacratus*, but differed from it in its much darker-brown color and in having a uniformly dark rather than bicolor tail. Allen [42] demoted *E. sacratus* to a subspecies of *E. caudatus*—an opinion with which Ellerman & Morrison-Scott [1] agreed. Hoffmann [7] examined many specimens and concluded that there were similarities in skull and cranial size between *E. c. caudatus*, *E. c. sacratus* and *E. c. umbrinus*. Wilson and Reeder [9,13] recognized these three taxa to be distinct species based on karyotypes and differences in skull size, while Motokawa and Lin [43] considered *E. sacratus* and *E. umbrinus* to be subspecies of *E. caudatus*. Motowaka *et al.* [44] considered the larger *E. caudatus* and smaller *E. sacratus* to be distinct species, and included three subspecies: *E. soluensis*, *E. umbrinus*, and *E. sacratus* as subspecies of *E. caudatus*. Wilson & Mittermeier [3] elevated *E. umbrinus* to full species without reason, while Wei *et al.* [45] regarded it to be a subspecies of *E. caudatus*. Our data support recognizing *E. sacratus* as a valid species, and recognizing *E. umbrinus* to be a subspecies of *E. caudatus*. In morphology, *E. sacratus* can be differentiated from *E. caudatus*, but *E. umbrinus* cannot. Our data do not support the opinion of Motowaka *et al.* [44].

Gruber [46] described *Episoriculus soluensis* and considered it a separate species. Abe [47] reviewed specimens collected in central Nepal and compared them with those from eastern Nepal by Gruber [46], and both suggested that *E. soluensis* was a subspecies of *E. caudatus*, but also possibly synonymous with *E. sacratus*. Hoffmann [7] treated *E. soluensis* as a synonym of *E. caudatus*—an opinion with which Wilson and Reeder [9,13] and Motokawa and Lin [43] agreed. Ohdachi *et al.* [48] regarded two samples from Nepal to be *E. caudatus soluensis* following Abe [47], and sequenced *CYTB*. Abramov *et al.* [2] then used these two sequences to reconstruct a system tree, and after determining that *E. soluensis* constituted a distinct clade from *E. caudatus*, advocated for them being treated as distinct species. In our tree, four samples from Yadong and Nyalam cluster with the two *E. soluensis* samples of Ohdachi *et al.* [48], and these four specimens are similar in having dark-brown ventral hair, and light-yellowish-brown dorsal hair. The tail length of our four specimens is longer than the head length, the skull parietal bone is relatively protruding, there are four upper single cusp teeth, the posterior cusp teeth of the maxillary incisor are similar in height to the first upper single cusp teeth, and the cusp teeth are light brown. These features are basically consistent with Gruber’s [46] original description, and the description of *E. soluensis* of Wilson & Mittermeier [3]. Accordingly, we regard *E. soluensis* to be a distinct species, and report it for the first time from China.

While Ellerman & Morrison-Scott [1] regarded *Episoriculus baileyi* to be a subspecies of *E. caudatus*, Abe [47, 49] identified the two to be morphologically distinct and sympatric in Nepal. Hoffmann [7] examined species from Burma and Nepal and considered *E. baileyi* to be a subspecies of *E. leucops*, an opinion with which Wilson and Reeder [9,13] agreed. Based on external and cranial morphology, Motokawa & Lin [44] re-evaluated the taxonomic status of *E. baileyi*, and considered it to be a valid species of *Episoriculus*. This species could be distinguished from other *Episoriculus* in the combination of its robust first upper incisor, long rostrum and upper unicuspid row, large tympanic ring, and high ascending ramus of the mandible—an opinion with which Wilson & Mittermeier [3] agreed. Because of a lack of specimens, we cannot investigate the status of *E. baileyi*. We provisionally follow Motokawa & Lin [44], but the taxonomic status of this species requires further investigation.

While Ellerman &Morrison-Scott [1] considered *E. fumidus* to be a subspecies of *E. caudatus*, Jameson and Jones [11] considered it to be a distinct species based on its geographical isolation and morphological divergence—an opinion with which Hoffmann [7], Wilson & Reeder [9,13], and Motokawa & Lin [44] agreed. Dubey *et al.* [23] inferred that *E. fumidus* (the only representative of the genus in their study) was a sister group of *Chodsigoa* with strong support in the *APOB* gene tree—an opinion with which He *et al*. [15] agreed. Based on the sequence of Dubey *et al.* [23] and He *et al*. [15], Abramov *et al.* [2] regarded *fumidus* do not belong in *Episoriculus*, and established the genus *Pseudosoriculus* for it—an opinion supported by our analyses.

We identify what appears to be a new cryptic species (*E*. sp.) from low-elevation areas in Motuo County, Tibet, which forms a separate branch in our system tree. However, with only two specimens available, we cannot accurately describe its morphology. Further specimens and molecular data are required to accurately resolve the taxonomic status of this taxon.

Our phylogenic analysis consistently roots *E. macrurus* as an individual lineage. This species, which has large genetic distance from congeners, in phylogenetic trees is usually located in the outermost or most basal part of the genus *Episoriculus.* It has the longest tail in the genus, and differs from congeners in skull and tooth morphology. For these reasons we speculate it retains some of the most primitive traits in genus *Episoriculus* or Nectogalini. We consider that *E. macrurus*, *E. caudatus*, *E. sacratus*, *E. soluensis*, *E. leucops*, *E. baileyi*, and a cryptic species *E.* sp. are all valid taxa. Both *E. baileyi* and *E.* sp. require additional specimens or molecular sequences to resolve their taxonomic status.

## Conclusion

Based on molecular and morphological analyses, the genus *Episoriculus* comprises at least six valid species: *E. baileyi*, *E. caudatus*, *E. leucops, E. macrurus, E. sacratus, E. soluensis,* and the potentially undescribed *E.* sp.

## Acknowledgments

This research was funded by the National Natural Science Foundation of China (32370496, 31970399). We are particularly grateful to Robert Murphy for the correct scientific questions. We are also grateful to Yingting Pu and Jiao Qing for their assistance with this study.

## Supporting information

**S1 Table** External and selected cranial measurements of *Episoriculus* species.

**S2 Table** Results of ABGD species definition based on *CYTB* gene.

**S3 Table** Posterior probabilities supporting three species (*Episoriculus caudatus*, *E. sacratus*, and *E. umbrinus*) as potential species using different algorithms and priors.

